# Ultraconserved elements occupy specific arenas of three-dimensional mammalian genome organization

**DOI:** 10.1101/235242

**Authors:** Ruth B. McCole, Jelena Erceg, Wren Saylor, Chao-ting Wu

## Abstract

This study explores the relationships between three-dimensional genome organization and the ultraconserved elements (UCEs), an enigmatic set of DNA elements that show very high DNA sequence conservation between vertebrate reference genomes. Examining both human and mouse genomes, we interrogate the relationship of UCEs to three features of chromosome organization derived from Hi-C studies. Firstly, we report that UCEs are enriched within contact ‘domains’ and, further, that the UCEs that fall into domains shared across diverse cell types are linked to kidney-related and neuronal processes. In ‘boundaries’, UCEs are generally depleted, with those that do overlap boundaries being overrepresented in exonic UCEs. Regarding loop anchors, UCEs are neither over- nor under-represented, with those present in loop anchors being enriched for splice sites compared to all UCEs. Finally, as all of the relationships we observed between UCEs and genomic features are conserved in the mouse genome, our findings suggest that UCEs contribute to interspecies conservation of genome organization and, thus, genome stability.

## INTRODUCTION

Chromosome organization in the mammalian nucleus is strikingly orchestrated, like a tightly arranged symphony played throughout the organism’s lifespan and composed by evolutionary forces. To explore this process of evolutionary ‘composition’, we are investigating the relationships between chromosome organization and sequence evolution in the mammalian genome, focusing our study on some of the most highly conserved regions – the ultraconserved elements (UCEs) (1-3). UCEs show staggering levels of interspecies sequence conservation, demonstrating perfect sequence identity extending ≥ 200 bp between species that diverged 90 to 300 million years ago and comprising one of the most mysterious findings of modern comparative genomics (4,5). While UCEs have been found to encompass a variety of functions, including enhancer, promoter, splicing, and repressive activities (1-3,6-23), these functions arguably fall short of explaining ultraconservation, *per se.* We have suggested that UCEs may maintain their sequence conservation through a mechanism involving the pairing and then comparison of allelic UCEs, followed by loss of fitness should mutations in UCEs or rearrangements that disrupt UCE pairing be detected (20,24-27). Such a mechanism would protect genome integrity in the body overall and, at the organismal level, promote ultraconservation over evolutionary timescales. Consistent with this model, UCEs are associated with regions of elevated synteny between different species (2,28-35). Furthermore, and in line with our proposal that disruptions of UCEs or UCE pairing lead to loss of fitness, the genomes of healthy individuals are generally not disrupted in the vicinity of UCEs (4,24,25,27), while this pattern does not hold for genomes representing the cancerous state (27), or individuals with neurodevelopmental disorders or mental delay and congenital anomalies (36). Highly conserved noncoding sequences have also been demonstrated to interact in three dimensions (37), adding weight to our proposal that interactions between UCEs in the nucleus may be important to their function. Finally, and of direct relevance to the proposal that allelic UCEs may pair, is the growing number of phenomena in a wide range of species, including mammals, demonstrating the capacity of somatic genomes to support localized or whole chromosome pairing (as reviewed by (38)).

Here, we examine UCEs in the context of the three-dimensional organization of the genome, considering in particular three features revealed by chromosome conformation capture (Hi-C) studies. The first two are contact ‘domains’ (also called topologically associated domains, or TADs) and ‘boundaries’; contact domains are regions displaying frequent intra-regional interactions, while ‘boundaries’, which flank contact domains, are characterized by a paucity of interactions that traverse them (39-47). A third type of interaction, at the edges of and within some domains, involves the three-dimensional association of ds-linked regions, wherein the intervening segment generates a loop, and the interacting regions, known as ‘loop anchors’, bring together regulatory regions, such as enhancers, silencers, and promoters (46). In concordance with the functional importance of these three features, global comparisons between human and mouse reveal that the positions of around half of domains, boundaries, and loops are shared (43,46) and, when shuffling does occur during the evolution of species, domains tend to be preserved as units (48). Thus, disrupting functional three-dimensional contacts inside domains may be disadvantageous, perhaps even oncogenic (49-53).

This study considers our proposal that ultraconservation protects genome integrity (24,25,27) and hypothesizes that UCEs contribute to the preservation of domains over evolutionary time. In particular, we predicted that UCEs would be enriched within domains. In line with this prediction, a recent publication reported that clusters of highly conserved noncoding elements (CNEs) correlate with the spans of domains encompassing genes involved in development (54); although the thresholds for the length and identity used in this publication to define CNEs (>50bp of 70-90% conservation between human and chicken genomes) are much less stringent than those used to define UCEs, the findings are intriguing in light of our proposal. To test our hypothesis, we examined 10 human and 6 mouse Hi-C datasets (43,46,55) and asked whether UCEs are enriched in or depleted from domains, boundaries, or loop anchors. Excitingly, UCEs proved to be significantly enriched in domains. Our results further revealed that the subset of UCEs occurring in domains that are present in multiple cell types is associated with kidney-related processes. In contrast, UCEs are generally depleted from boundaries and neither enriched nor depleted from loop anchors. The UCEs that do, nevertheless, occur in boundaries and loop anchors are predominantly exonic, with those in loop anchors enriched in splice sites. Our findings demonstrate that UCEs show specific, conserved relationships to domains, boundaries, and loops, hinting that UCEs may play a role in establishing and maintaining genomic organization.

## MATERIALS AND METHODS

### Hi-C data sources

The genomic coordinates for domains, boundaries, and loop anchors were obtained from the published Hi-C datasets (Supplementary Tables S1 and S2) (43,46,55). When required the coordinates were converted using UCSC Genome Browser tool liftOver (http://genome.ucsc.edu/cgi-bin/hgLiftOver) to hg19 genome assembly. To avoid counting a same region multiple times, overlapping genomic coordinates were merged providing a final list of coordinates that may differ from originally reported ones in the respective publications. For each individual and pooled Hi-C dataset after coordinate merging the information about the number of regions, median size (bp), coverage (bp), and proportion of covered genome (%) was reported (Supplementary Tables S1 and S2).

To account for data variability given that the resolution between datasets varied based on the amount of biological material, applied Hi-C protocol (46,56-58), and sequencing depth, each chromosomal feature (domain, boundary, loop anchor) was inspected individually within a single dataset, in addition to pooling across multiple datasets.

### UCE data sources

The UCEs encompass a dataset representing 896 HMR-HDM-HC elements as previously reported in hg18 genome assembly (24,27). All UCE genomic coordinates were lifted over to hg19 genome assembly and made available in Supplementary Table S3A.

To obtain the respective UCE coordinates in mm9 genome assembly, a read aligner tool bowtie2 was used (59). To start, the UCEs sequences in fasta format were converted to fastq format, and then subsequently mapped using bowtie2 with the following parameters: ‘bowtie2 -p 4 -x /bowtie_index/mm9 --very-sensitive -t -S UCEs_mm9.sam -U sequences_hg18_allUCEs.fastq’. The end-to-end alignment was chosen to avoid ‘trimming’ or ‘clipping’ of some read characters from ends of the alignment. The output file in SAM format was first converted to a sorted BAM format (samtools view -bS UCEs_mm9.sam I samtools sort - UCEs_mm9_sorted) using SAMtools (60), and then to a BED file (‘bedtools bamtobed −i UCEs_mm9_sorted.bam > UCEs_mm9_sorted.bed’) using BEDTools (61). Upon filtering matches in the mouse genome that were <200 bp (UCE_153, UCE_600) or differed more than 15 bp in length to hg19 coordinates (UCE_733), we obtained 893 UCEs in mm9 genome assembly. The output mm9 UCE coordinates adjusted to 1-based system were reported in Table S3B. Three UCEs were not recovered in the mouse genome, namely UCE_153, UCE_600, and UCE_733. The first two omitted UCEs spanned less < 200 bp in the mouse genome, and thus did not satisfy our initial UCE definition that required a UCE to be ≥ 200 bp in length. The third UCE_733 was excluded since it differed more than 15 bp in length to the human counterpart, and it falls within a mouse intergenic region as opposed to its human UCE sequence that lies within the intron of POU6F2 gene.

During analysis UCEs were further subclassified into exonic, intronic or intergenic elements as previously described (24,27) to allow for finer dissection of UCE relation to chromosome organization (Table S3).

### Data sources for correlation analysis

The CNV/CNA data previously assembled (27) was inspected in healthy individuals representing classical inherited CNVs (62-69), and CNAs derived from 52 different cancers (70-86). All CNV and CNA genomic coordinates were lifted over to hg19 assembly and collapsed.

The genomic coordinates for hg19 assembly were downloaded from the UCSC Table Browser (87), coordinates for genes, introns, and exons were derived from group - Genes and Gene Predictions, track – UCSC genes; repetitive element were from group – Repeats, track - RepeatMasker, and contain SINE, LINE, and LTR repetitive elements; segmental duplications were from group- Repeats, track-Segmental Dups, CpG islands were from group – Regulation, track - CpG Islands, and open chromatin identified by the ENCODE project (88) from group – Regulation, track – Open Chrom Synth for cell lines that match Hi-C datasets (GM12878, H1-hESC, K562, HeLa, HeLa-Ifna4h, HUVEC, NHEK).

### Depletion or enrichment analysis of UCEs in specific genomic regions

The enrichment or depletion of UCEs in genomic regions of interest such as domains, boundaries, and loop anchors was assessed using established methods previously reported in our publications (24,25,27). Briefly, observed overlap between for instance UCEs and domains were compared to mean expected overlap, which was produced by placing randomized set of elements that match UCEs in number and length 1,000 times in the genome. The distribution of expected overlaps was assessed for normality using Kolomogorov-Smirnov (KS) test. In a case normality was observed, Z-test comparison between observed and expected overlaps was reported, which when significant indicated depletion if ratio between observed and mean expected overlaps (obs/exp) was below 1.0, or enrichment if the obs/exp ratio was above 1.0. In instances where normality was not observed, the proportion of expected overlaps equal to, or more extreme than the observed overlap together with obs/exp ratio was reported. During further dissection of relation between UCEs, which were classified to intergenic, intronic, and exonic elements, to genomic regions of interest random set of elements used to calculate mean expected overlap was pooled from the matching genomic regions, *i.e.* only intergenic, intronic, or exonic ones.

Distribution of expected overlaps and observed overlap (colored line) were visualized using histograms, where corresponding statistical values were reported based on the results of the KS test.

### Correlation analyses

The genome was divided into bins of equal sizes. Within each bin, the fraction of sequence occupied by each control feature was calculated, as was that of UCEs, except in the case of GC content, where it was calculated as the fraction of G + C. Then genome-wide correlations within each bin were preformed among feature densities or GC content. The Spearman correlation coefficients and matching P-values were provided in two flavors, either for a pairwise comparison between two features, or as a part of partial correlation approach, which assesses whether the correlation between two features was affected by co-correlation with the third genomic feature. The obtained Spearman correlation coefficients were visualized as a heatmap.

### Distribution of intergenic, intronic, and exonic UCEs that overlap domains, boundaries, and loop anchors

Analysis on the distribution of intergenic, intronic, and exonic UCEs (reported from depletion/enrichment analysis in Supplementary Table S4A) was performed by comparing a control set (all 896 UCEs from Supplementary Table S3A) to UCEs that overlap feature of interest, *i.e.* either domains, boundaries, or loop anchors (Supplementary Table S1). A statistical significance was determined using chi-squared test, and reported for pooled (Supplementary Table S5A), and other human domain datasets from other cell lines (Supplementary Table S5B).

To determine the amount of domains, boundaries, and loop anchors that fall within intergenic, intronic, and exonic regions, the overlap between two features (*i.e.* intergenic regions in domains) was calculated using bedtools intersect (61). The information on an overlap such as number of intervals, median interval size (bp), coverage (bp), percentage of genome, and percentage of a Hi-C feature coverage is reported for Pooled domains, boundaries, and loop anchors (Supplementary Table S5C).

### Gene ontology

Functional association of UCEs to the gene ontology (GO) terms of nearby genes was determined against the full set of 896 human (or 893 mouse) UCEs as a background using the Genomic Regions Enrichment of Annotations Tool (GREAT) (89). The background provided a control that observed functional association for domain invariant UCEs and exonic UCEs that overlap boundaries and loop anchors was inherent to those UCE subsets, and not the full set of UCEs.

### Analysis on the number of times each UCE is overlapped by the individual domain dataset

Each assessment of overlap between every UCE and every individual domain dataset is assigned a score of 1 (intersect) or 0 (no intersect). The summation of scores across all individual datasets resulted in a total score for each UCEs. Those UCEs with the highest score, *i.e.* that are confirmed by all individual datasets to overlap domains, were termed domain invariant UCEs, and reported for human (Supplementary Table S7A), and mouse (Table S7B). Overlap between domain invariant UCEs from human and mouse was performed using UCE ID identifiers. Shared domain invariant UCEs between human and mouse, which are classified as exonic or intronic, were reported together with the assigned gene (UCSC) and gene abbreviation (NCBI) in Supplementary Table S7C.

## RESULTS

### UCEs are enriched within domains, depleted from boundaries, and indifferent to loop anchors

We began our studies by delineating how the Hi-C annotated genomic features of domains, boundaries, and loop anchors are related to the positioning of UCEs. To do this, we first collected published Hi-C datasets derived from nine human (Supplementary Table S1) and five mouse (Supplementary Table S2) tissues, representing a variety of cell types (43,46,55). As Hi-C annotated regions vary between studies due to differences in cell type, species examined, amount of starting material, Hi-C protocol (in-solution (43,55) or in-nucleus (46)), and sequencing depth, we examined each dataset individually in addition to querying datasets generated by combining datasets according to species and genomic feature (Supplementary Tables S1 and S2) (43,46,55). Regarding UCEs, our analyses used our previously defined dataset (Table S3), which comprises 896 elements that are ≥ 200 bp in length and identical in sequence within at least one of three groups of reference genomes (24,27). The three groups consist of the reference genomes of human, mouse, and rat (HMR), of human, dog, and mouse (HDM), and of human and chicken (HC), with the combined dataset of 896 UCEs designated as HMR-HDM-HC (Supplementary Table S3A). To obtain UCE positions in the mouse genome, we used the Bowtie 2 short read alignment program and recovered 893 orthologs (Materials and Methods; Supplementary Table S3B). UCEs were also subdivided into exonic, intronic, and intergenic categories, which were then examined jointly and separately for enrichment or depletion within the Hi-C annotations (Experimental Procedures). Of note, a UCE is considered exonic if any part overlaps an exon, hence, exonic UCEs may overlap splice sites and contain intronic sequence.

To assess whether UCEs are significantly enriched in or depleted from domains, boundaries, and loop anchors, we used our previously established method (24,25,27) (Figure 1), which compares the observed overlaps, in base pairs, between UCEs and Hi-C annotated regions to overlaps between a set of regions matched to UCEs in terms of number and length, but randomly positioned in the genome, calling the latter ‘expected overlaps’. Expected overlaps are generated 1,000 times to produce a distribution of expected overlaps, which is then tested for normality using the Kolomogorov-Smirnov (KS) test, and where the test affirms normality, a Z-test is used to compare the observed overlap with the distribution of expected overlaps to obtain a P-value describing the significance of any deviation. In cases where normality is not observed, the proportion of expected overlaps equal to, or more extreme than, the observed overlap is reported. In all cases, we report the ratio of observed to mean expected overlap (obs/exp). These steps ensure a tailored approach for each Hi-C dataset, enabling the comparison of datasets that differ in number of identified regions, median region size, and percentage of genome covered.

**Figure 1.**
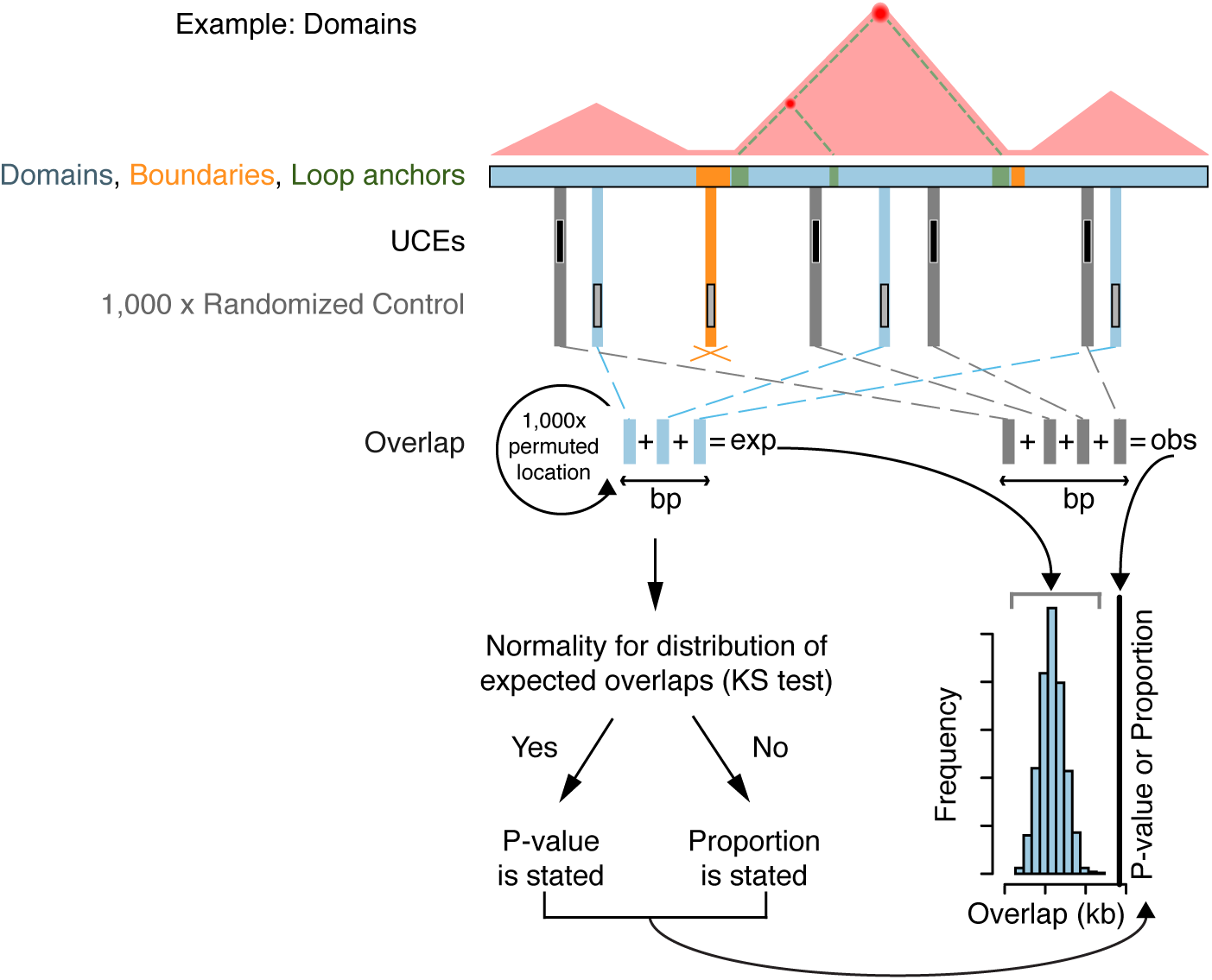
Strategy for assessing the relationship between UCEs and domains, boundaries, and loop anchors. We assess the relationship of UCEs (black) to domains (blue), boundaries (orange), and loop anchors (green) via a multi-step process, illustrated here with respect to domains. Throughout this and other figures, blue, orange, and green refer to analyses related to domains, boundaries, and loop anchors respectively. First, overlaps between UCEs and all domains in a dataset are summed to produce the observed overlap; as this example concerns domains, overlap between UCEs and boundaries are not tallied (orange cross). The observed overlap is then compared to a distribution of expected overlaps generated from the overlap of domains with each of 1,000 sets of control genomic sequences, matched to UCEs in number and length and randomly positioned in the genome. Finally, the distribution of the resulting 1,000 control overlaps is tested for normality using the Kolomogorov-Smirnov (KS) test and, when normality is observed, a P-value is reported to describe the significance of the deviation of the observed overlap from the distribution of expected overlaps. If normality is not observed, the proportion of expected overlaps equal to, or more extreme than, the observed overlap is stated.

We first analyzed ten datasets of domains, drawn from Dixon *et al.* (22) and Rao *et al.* (29), that examined nine human cell lines, whose origins spanned embryonic (hESC) and fetal (IMR90 lung fibroblast) development, cancer (HeLa, K562, and KBM7), and differentiated tissues (GM12878, HMEC, HUVEC, and NKEK), with IMR90 studied by both Dixon *et al.* and Rao *et al.* and thus contributing two datasets (Supplementary Table S1). The domains described by these datasets range in coverage from 83.18% of the genome for hESC domains from Dixon *et al.* (22) to 40.07% for HMEC domains from Rao *et al.* (29). Excitingly, we observed significant enrichment for UCEs within domains in eight out of ten datasets (4.22 × 10^-15^ ≤ P ≤ 0.020, 1.061 ≤ obs/exp ≤ 1.167; Supplementary Table S4A); the two in which enrichment was not seen represented HMEC and NHEK cells (29) from Rao *et al.* (Supplementary Table S4A). Combining all ten datasets, which included merging overlapping regions, produced a dataset, called ‘Pooled domains’, containing 293 regions covering 89.1% of the genome (Supplementary Table S1) that is also significantly enriched for UCEs (P = 2.77×10^-6^, obs/exp = 1.025; Figure 2A, Supplementary Table S4A). These results show that UCEs are overrepresented within Hi-C domains across many cell types, supporting the idea that there is an interrelationship between UCEs and three-dimensional chromosome conformation.

**Figure 2.**
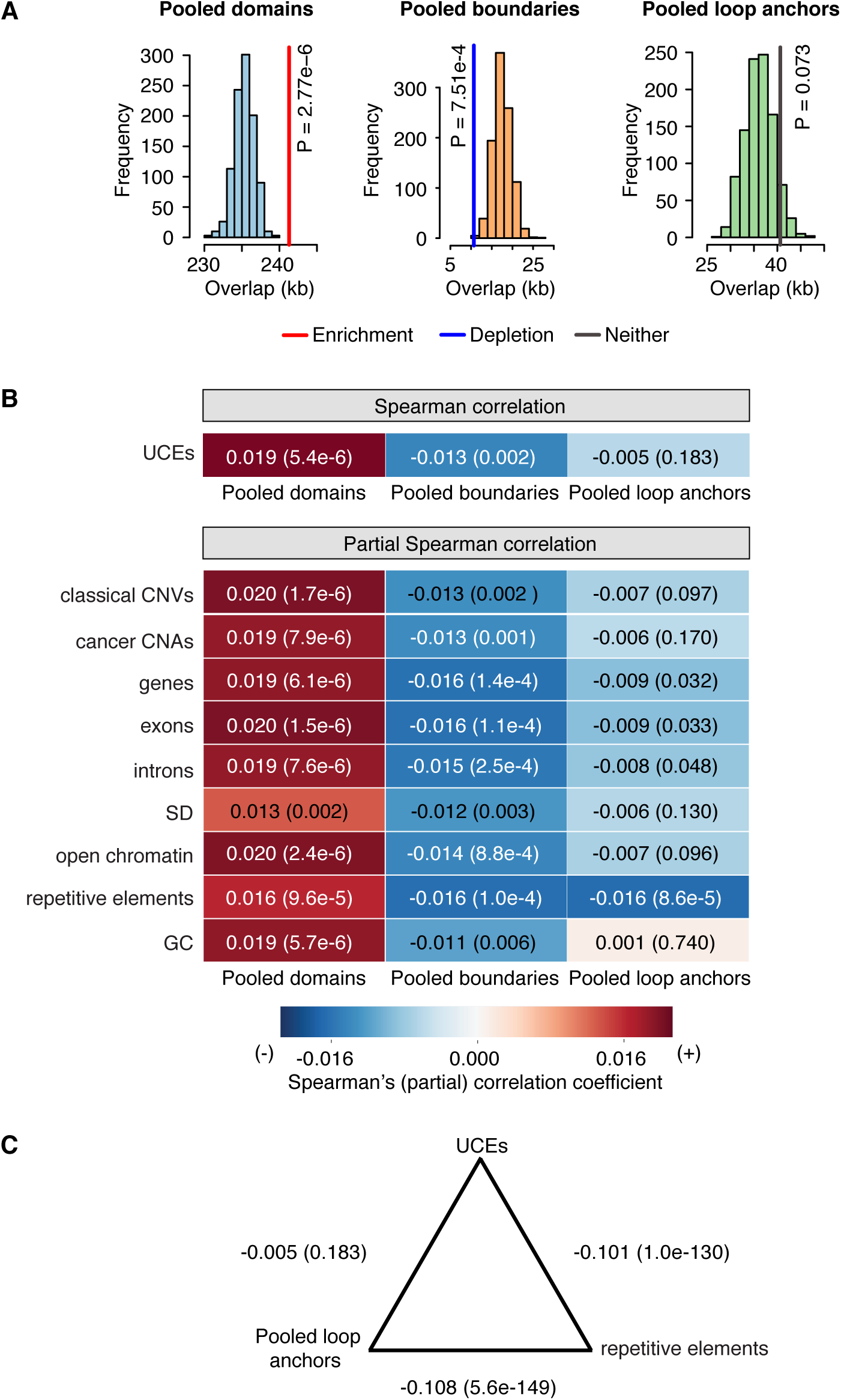
UCEs are enriched in Pooled domains, depleted from Pooled boundaries, and indifferent to Pooled loop anchors. **(A)** In the case of Pooled domains, the observed overlap (colored vertical line) of UCEs is significantly greater than the expected overlaps (red line; P=2.76×10^-6^, obs/exp=1.025). For Pooled boundaries, the observed overlap is significantly less than expectation (blue line; P=7.51×10^-4^, obs/exp=0.609). Observed overlap between UCEs and Pooled loop anchors does not deviate significantly from expectation (grey line; P=0.073, obs/exp=1.124). **(B)** Correlation analyses. Spearman correlation: Using pairwise Spearman correlation and splitting the genome into 50kb bins, the representation of UCEs is positively correlated with that of Pooled domains (P = 5.4×10^-6^), negatively correlated with that of Pooled boundaries (P = 0.002), and not significantly correlated with that of Pooled loop anchors (P = 0.183). Partial Spearman correlation: The positive and negative correlations between the positions of UCEs and Pooled domains (first column), and negative correlation between the positions of UCEs and Pooled boundaries (second column) remain significant even after accounting for the correlation between the positions of UCEs and nine control genomic features. The representation of UCEs and Pooled loop anchors (third column) is not significantly positively nor negatively correlated except when controlling for repetitive elements, explored in Panel C. **(C)** Although UCEs and Pooled loop anchors are not significantly correlated with each other (P = 0.183), pairwise correlation analyses of both UCEs and Pooled loop anchors show a highly significant negative correlation with repetitive elements (P=1.0×10^-130^ and P=5.6×10^-149^ respectively). **(B and C)**: Spearman (partial) correlation coefficients are reported in each box and by a heatmap; P-values are reported in parentheses.

We then examined datasets of boundaries from Dixon *et al.* (22). These datasets, which represent hESC and IMR90 cells and cover 3.95% and 3.58% of the genome, respectively (Supplementary Table S1), are significantly depleted of UCEs (respectively, P = 0.002, obs/exp = 0.516 and P = 0.025, obs/exp = 0.669; Supplementary Table S4A, although for IMR90, the P-value hovers at our significance cutoff). Merging the two datasets created a ‘Pooled boundary’ dataset, containing 3,715 regions and covering 6.59% of the genome (Supplementary Table S1), that is also depleted for UCEs (P=7.51×10^-4^, obs/exp 0.609; Figure 2A, Supplementary Table S4A). These findings reinforce our observation that UCEs do not commonly occur within Hi-C boundaries, and complement our previous observation that UCEs preferentially occur within domains.

Our next analysis concerned eight datasets of loop anchors provided by Rao *et al.* and representing GM12878, HeLa, HMEC, HUVEC, IMR90, K562, KBM7, and NHEK cells, with genome coverage ranging from 2.28 to 5.9% (Supplementary Table S4A). For all but two datasets, UCEs are neither enriched nor depleted (0.006 ≤ P ≤ 0. 480, 0.710 ≤ obs/exp ≤ 1.334; Supplementary Table S4A). Merging all eight datasets produced a dataset of ‘Pooled loop anchors’, comprising 18,331 regions and covering 13.58% of the genome (Supplementary Table S1), that is also neither enriched nor depleted for UCEs (P = 0.073, obs/exp 1.124; Figure 2A, Supplementary Table S4A). The overall lack of UCE enrichment in loop anchors is surprising, since many UCEs show enhancer-like properties (3,8,11,13,16-19,21), and enhancer-promoter interactions have been proposed to generate loops (46). Indeed, we did observe enrichment of UCEs in two of the eight datasets, HUVEC (P = 0.020, obs/exp = 1.322; Supplementary Table S4A) and NHEK (P = 0.006, obs/exp = 1.334; Supplementary Table S4A), suggesting that UCEs might be particularly involved in loop anchors in endothelial and epidermal cell types (HUVEC and NHEK cells, respectively).

### Relationships of UCEs to Hi-C annotations are robust

Having revealed positional relationships between UCEs and domains, boundaries, and loop anchors, we examined whether these relationships are robust to co-correlation with nine other genomic features. These features, which can be considered controls, included six that had previously shown to be non-randomly associated with UCE positions: copy number variants (CNVs), cancer-specific copy number alterations (CNAs), genes, exons, introns, and segmental duplications (SDs) (24,25,27). They also included open chromatin, since UCEs have been linked to transcriptional activity (1-3,6-9,11,13,15-23,90-94), repetitive elements, which UCEs avoid (1,24,25,27), and GC content, which is associated with the positions of CNVs (95). We divided the genome into equally sized bins and, because domains and the nine control features span a vast range of sizes, our analyses involved multiple iterations using a range of bin sizes (20, 50, and 100kb). Within each bin, the fraction of sequence occupied by each control feature was calculated, as was that of UCEs, except in the case of GC content, where it was calculated as the fraction of G + C (Materials and Methods). Genome-wide correlations were then determined with respect to each control within each bin.

Using pairwise Spearman correlation coefficients and associated P-values for the strength of correlation, we first determined that UCEs are significantly and positively associated with Pooled domains (P = 5.4×10^-6^; Figure 2B), significantly negatively correlated with Pooled boundaries (P = 0.002; Figure 2B) and not correlated with Pooled loop anchors (P = 0.183; Figure 2B). These results correspond well to UCE enrichment, depletion, and neither enrichment in nor depletion from Pooled domains, boundaries, and loops, respectively (Figure 2A). Then, using a partial correlation approach, we asked whether these correlations, or lack thereof, are influenced by co-correlation with any of the nine control genomic features. With a bin size of 50kb, the correlation between UCEs and Pooled domains remains significantly positive in all cases, indicating that it is robust to contributions from the control features (Figure 2B). Similarly, the negative correlation between UCEs and Pooled boundaries remains robust to all control features (Figure 2B). As for Pooled loop anchors, the correlation with UCEs is insignificant in all cases but one, consistent with UCEs being neither enriched nor depleted in Pooled loop anchors (Figure 2B). The one exception pertains to repetitive elements, where the correlation is significantly negative. Investigating this further, we discovered a negative correlation between UCEs and repetitive elements (P = 1.0×10^-130^; Figure 2C), which is unsurprising, as UCEs are non-repetitive (1,24,25) and avoid insertions of repetitive elements (96). We also uncovered a strong negative correlation between Pooled loop anchors and repetitive elements (P = 5.6×10^-149^; Figure 2C). Thus, it may be that, while a significant negative correlation exists between UCEs and Pooled loop anchors, it is secondary to the strong negative correlation between repetitive elements and both UCEs and Pooled loop anchors. Altering the sizes of the genomic bins to 20kb (Supplementary Figure S1A) and 100kb (Supplementary Figure S1B) produced very similar results. We note that with all three bin sizes, when controlling for GC content, correlation between UCEs and Pooled loop anchors is positive, but that this is significant only in the case of the 20kb bin size (P = 0.011; Supplementary Figure S1). Taken together, the positioning of UCEs relative to domains, boundaries, and loop anchors is robust to co-correlation with nine other genomic features.

### Positioning of UCEs within Hi-C annotations is conserved between human and mouse

Since UCEs are defined by their extreme evolutionary conservation between species, we went on to ask whether the relationships observed between UCEs and domains, boundaries, and loop anchors in the human genome are conserved in the mouse genome. Accordingly, we turned to the 893 mouse orthologs (Supplementary Table S3B) of our human UCEs and three Hi-C studies (43,46,55) addressing mouse embryonic stem cells (mESC), blood (B-lymphoblasts), neuronal precursor cells (NPC), post-mitotic neurons, and cortical tissue (Supplementary Table S2). We found that the relationships of UCEs to domains, boundaries, and loop anchors are evolutionarily conserved. For domains, we examined six datasets covering between 29.02% of the genome, in the case of lymphoblasts from Rao *et al.,* and 92.49%, in the case of neurons from Fraser *et al.* (Supplementary Table S2). All six datasets were significantly enriched for UCEs (5.52×10^-10^≤ P ≤ 0.002; 1.020 ≤ obs/exp ≤ 1.260; Supplementary Figure S2, Supplementary Table S4B). For boundaries, we examined a dataset that is described by Dixon *et al.* (22) to be common to both mESC and cortex tissue and covering 8.11% of the genome, calling this dataset ‘Mouse common boundaries’ (Supplementary Table S2). This dataset shows significant depletion for UCEs (P = 4.68×10^-7^, obs/exp = 0.452; Supplementary Figure S2, Supplementary Table S4B). Finally, with respect to loop anchors, we examined one dataset from Rao *et al.* from lymphocytes, which covers 2.16% of the genome (Supplementary Table S2) and is neither enriched in nor depleted of UCEs (P = 0.090, obs/exp = 0.701; Supplementary Figure S2, Supplementary Table S4B).

### When UCEs are found in boundaries and loop anchors, they show an excess of exonic UCEs associated with RNA processing

Having established that UCEs are differentially associated with domains, boundaries, and loop anchors, we queried whether specific subsets of UCEs might be driving the associations. In particular, we examined intergenic, intronic, and exonic UCEs separately, since these subdivisions have behaved distinctly in our previous studies. For example, the extent to which UCEs are depleted from CNVs, and the depth of the plunge in AT content at UCE boundaries are more extreme for intergenic and intronic UCEs than for exonic UCEs (24,25,27). First, we examined all the individual datasets for domains as well as Pooled domains and found that, in all cases, there is no significant deviation from expected in the observed proportions of intergenic, intronic, and exonic UCEs (0.131 ≤ P ≤ 0.892; Figure 3A, Supplementary Tables S5A and S5B). Note that, Pooled domains includes all 896 UCEs, including boundary UCEs, because the boundaries of some cell types are organized as domains in other cell types. Thus, the proportions of UCEs in Pooled domains are the same as those within the entire UCE dataset.

**Figure 3.**
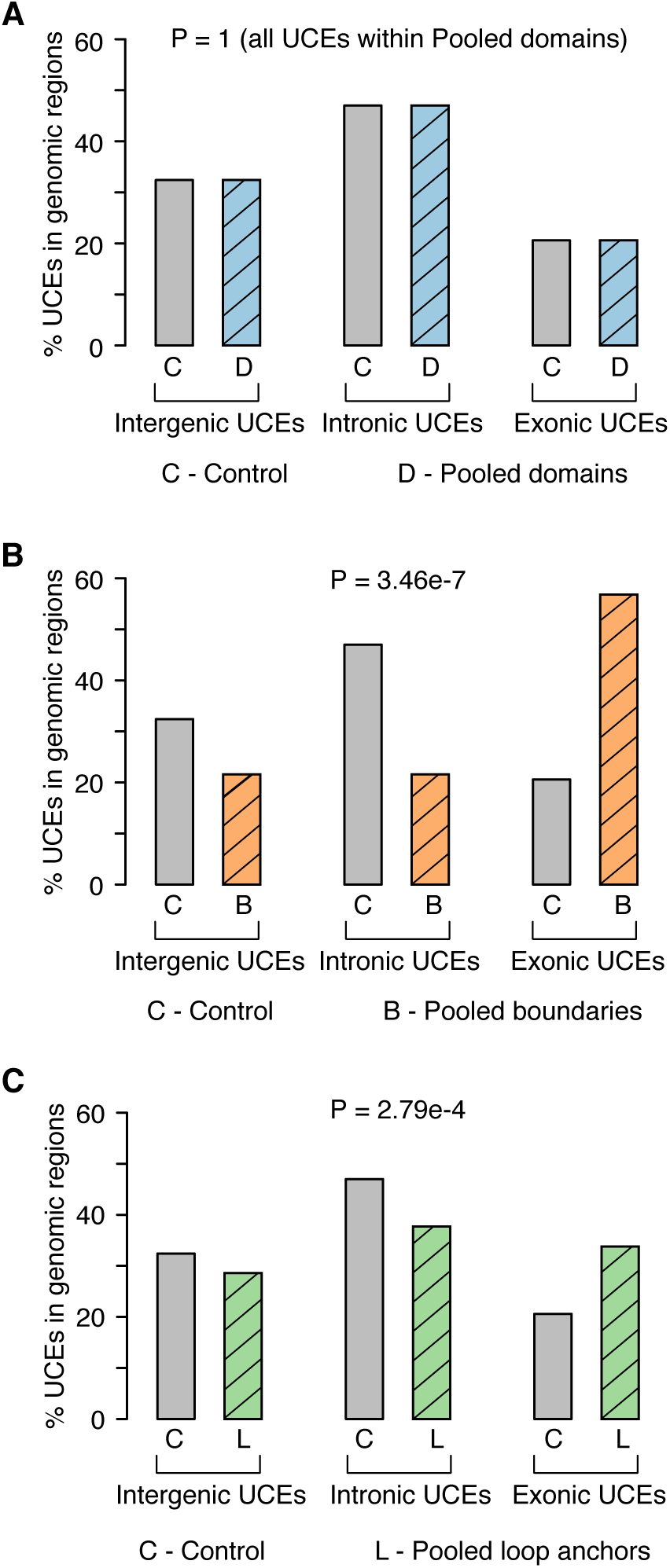
Underrepresentation of intronic and intergenic UCEs in Pooled boundaries and loop anchors are accompanied by overrepresentation of exonic UCEs. Proportions of intergenic, intronic, and exonic UCEs that overlap Pooled domains (**A**, blue), boundaries (**B**, orange), and loop anchors (**C**, green) compared to the full set of 896 UCEs as a control (grey). Pooled domains that are not significantly different compared to the control set since **(A)** All UCEs fall within pooled domains, so no P-value is calculated. **(B-C)** Pooled boundaries (**B**; chi-squared test, P = 3.46×10^-7^) and Pooled loop anchors (**C**; chi-squared test, P = 2.79×10^-4^) both show a significant overrepresentation of exonic UCEs and an under-representation of intronic and intergenic UCEs, as compared with the full UCE set.

For Pooled boundaries, the distribution of intergenic, intronic, and exonic UCEs deviates significantly from that of the full set of UCEs (P = 3.46×10^-7^; Figure 3B, Supplementary Table S5A). We found a depletion of intergenic and intronic UCEs, with 21.6% (8 out of 37) and 21.6% (8 out of 37), respectively, in boundaries, as compared to the expected 32.4% (290 out of 896) and 47.0%, respectively. In contrast, exonic UCEs are overrepresented, with 56.8% (21 out of 37) in Pooled boundaries, while making up only 20.6% of all UCEs. The overrepresentation of exonic UCEs is even more striking in light of the fact that the majority (52.7%) of Pooled boundary DNA is intronic (109Mb), with only a small fraction (5.3%) being exonic (11Mb) (P = 2.89×10^-44^; Materials and Methods; Supplementary Table S5C).

We also found significant deviation of the proportions of intergenic, intronic, and exonic UCEs in Pooled loop anchors (P = 2.79×10^-4^; Figure 3C; Supplementary Table S5A). Intergenic and intronic UCEs represent only 28.6% (44 out of 154) and 37.7% (58 out of 154) of UCEs, respectively, whereas 32.4% and 47.0% of the full UCE set are intergenic and intronic, respectively. As in Pooled boundaries, exonic UCEs are overrepresented at 33.8% (52 out of 154) as compared to 20.6% of all UCEs. These proportions deviate significantly from expected based on the sequence composition of Pooled loop anchors, which is 47.0% intronic and only 5.02% exonic (P = 5.38×10^-60^; Supplementary Table S5C). These results point to intronic, and, to some extent, intergenic, UCEs as drivers of depletion from Pooled boundaries and to exonic UCEs as the dominant type of UCE within both Pooled boundaries and loop anchors.

We next used the Genomic Regions Enrichment of Annotations Tool (GREAT) (47) and discovered that exonic UCEs in Pooled boundaries and Pooled loop anchors are enriched for GO terms associated with RNA processing (Supplementary Figures S3A and S3B), and this is in line with previous reports that exonic UCEs are associated with RNA processing, including splicing (1,12,14,51,97-100). Considering further the structure of exonic UCEs, themselves, we found that 76% (16 out of 21; Supplementary Table S6A) and 82% (43 out of 52; Supplementary Table S6B) of exonic UCEs in Pooled boundaries and loop anchors, respectively, partially overlap introns and hence cover splice sites, as compared to 57% in the full set of exonic UCEs (107 out of 185; Supplementary Table 3C). Thus, while exonic UCEs in Pooled boundaries are not enriched for splice sites (P= 0.07; Supplementary Table S6A), those in Pooled loop anchors are (P = 1.82×10^-4^; Supplementary Table S6B). These results suggest a two-layered association of UCEs with RNA processing, whereby UCEs are associated with genes involved in RNA processing and UCEs may also help the splicing of these very same genes. This double association may be of particular importance to the presence of UCEs in loop anchors, and raises the possibility that loop anchors containing UCEs may exist to assist in particular splicing mechanisms.

### UCEs within domains that are shared in many cell types are associated with kidney-related processes

While domains vary between cell types, many are nevertheless shared and, thus, we next focused on UCEs that occur within domains common across multiple cell types; these might address the functional significance underlying the enrichment of UCEs. We first identified 124 UCEs that overlap domains in all 10 individual human datasets across diverse cell types (Supplementary Table S1), calling these ‘human invariant domain UCEs’ (Supplementary Table S7A), as well as 310 UCEs that overlap domains identified by all 6 mouse datasets (Supplementary Table S2), calling these ‘mouse invariant domain UCEs’ (Supplementary Table S7B). Using GREAT, these human and mouse invariant domain UCEs were compared to the full UCE sets in humans and mouse, respectively, revealing an association with kidney-related GO terms for human invariant domain UCEs (Supplementary Figure S4A, asterisked terms). Terms related to kidney biology were also obtained in the case of mouse UCEs although, here, other terms were attained as well, some with greater significance (Supplementary Figure S4B, asterisked terms). These findings are corroborated by the association with kidney-related processes, as well as neuronal development, of the 74 UCEs shared between the human and mouse invariant domain UCE datasets (Supplementary Figure S5 and Supplementary Table S7C). The proportions of invariant domain UCEs assigned as intergenic, intronic, and exonic do not significantly deviate from proportions of the entire UCE set (control), except for the mouse invariant domain UCEs, where we saw an overrepresentation of intronic and exonic UCEs, with intergenic UCEs underrepresented (P = 8.51×10^-4^; Supplementary Table S7D). Of note, Ahituv *et al.* (101) reported two mice out of a total of 102 individuals with a homozygous deletion for UCE 329 (our invariant domain UCE 159) to have only one kidney, the population frequency among humans for this condition being estimated at 1 in 1,000 live births. In brief, functions related to kidney development might be a feature of UCEs within domains shared among diverse cell types.

## DISCUSSION

Our findings reveal a non-random UCE distribution among three main arenas of three-dimensional genome organization, with UCEs being enriched in domains, depleted from boundaries, and indifferent to loop anchors (Figure 4). The UCEs that do occupy boundaries and loop anchors display an overrepresentation of exonic UCEs, and in loop anchors, those UCEs are enriched for overlap with splice sites, suggesting a specific involvement of loop anchors containing UCEs in splicing. With respect to UCEs in domains that do not vary between cell types, they are, as a group, significantly associated with kidney-related and neuronal gene ontologies.

**Figure 4.**
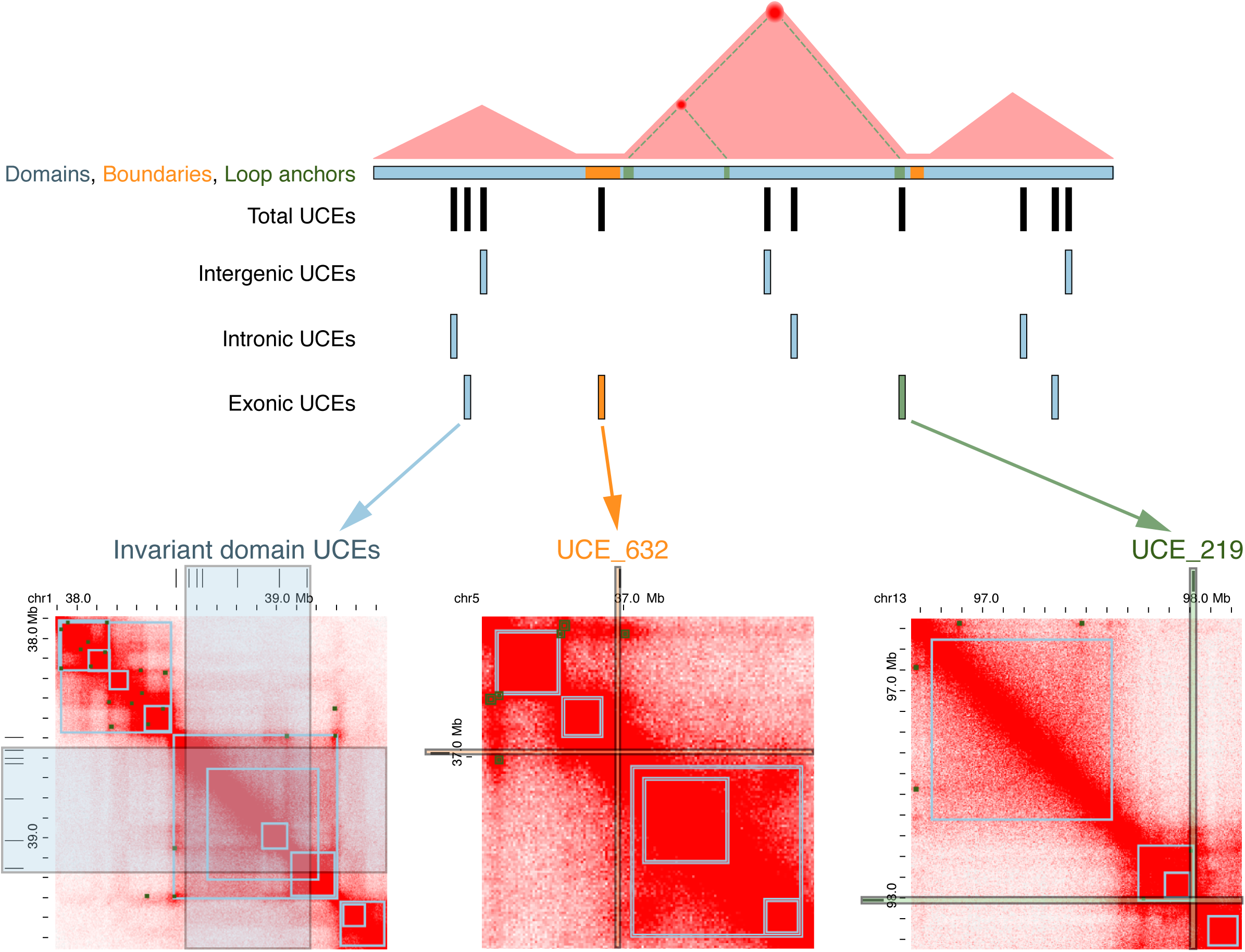
Schematic representation summarizing the relationship between chromosome organization and UCEs. *Top:* Domains (blue) are enriched in UCEs, boundaries (orange) are depleted, and loop anchors (green) are neither enriched nor depleted. *Bottom:* Examples of UCEs in each of the three genomic features as defined by Hi-C annotation of human GM12878 cells (46) using the Juicebox tool (109). Left, invariant domain UCEs; middle, UCE in a boundary; right, UCE in a loop anchor. Domain and loop anchor calls (squares in grey outline) are indicated on the heatmaps as available in Juicebox (109). UCEs are depicted in numbers not representative of their true occupancies.

These findings tying UCEs to genome organization are especially intriguing in light of the proposal that UCEs may contribute to genome integrity through yet another potent organizational feature of genomes - allelic and homolog pairing (20,24-27). Indeed, they raise the question of whether UCEs contribute to the establishment of domains, and/or whether the evolution of a domain promotes the fixation of UCEs within the domain. Consistent with this, Harmston *et al.* (54) recently reported that clusters of CNEs predict the span of domains, suggesting that CNEs might be involved in chromatin folding. For example, since some UCEs embody enhancer activity (1-3,6-9,11,13,15-23) and, thus, are likely to participate in enhancer-promoter interactions, might that activity help define chromosomal contacts? Separately, but not exclusively, might selection against changes that disrupt chromosomal domains promote sequence invariance and, thus, ultraconservation? Specifically, if, as we have proposed (24,25,27), rearrangements that disrupt the pairing of allelic UCEs are culled, then UCEs will contribute to the structural invariance of genomic regions in which they lie. In this way, UCEs may have enhanced the capacity of certain regions to evolve the intra-regional contacts that, today, define contact domains.

The strong association of invariant domain UCEs with kidney-related and neuronal GO categories was intriguing and merits further exploration. In this light, it may be noteworthy that evolution of the kidney has been argued to be an early defining process in the emergence of vertebrates (102). If so, that evolution may have benefitted from the genome stability provided by UCEs. Finally, we mention that homolog pairing in human kidney cells along an entire chromosome arm has been associated with renal oncocytoma (103), an observation that may ultimately prove relevant to our finding that UCEs in invariant domains are associated with kidney-related GO terms.

Our studies have also shown that, while boundaries are generally depleted of UCEs, 21 of the 37 UCEs found in boundaries are exonic, constituting an enrichment of exonic UCEs in boundaries. Of the 21 boundary exonic UCEs, two (UCEs 632 and 633) are in the *Nipbl* gene, which is a cohesin loading factor that, when mutated, leads to a developmental disorder known as Cornelia de Lange syndrome (104). Given that cohesin binding is implicated in sister chromatid cohesion and gene expression (105,106), ultraconservation within *Nipbl* may speak to this gene’s importance in genome structure and function. Indeed, a recent study demonstrated that depletion of *Nipbl* in mouse affects reorganization of chromosome folding (107). Furthermore, the evolutionarily conserved position of the *Nipbl* gene within boundaries may suggest that the lack of three-dimensional associations across a boundary may also be important for its expression.

Turning to loop anchors, their lack of enrichment in UCEs chimes with other findings arguing that loops are evolutionarily dynamic (32). Their dynamic nature is consistent with the malleability of enhancers over evolutionary time and thus, also, of enhancer-promoter interactions, both of which make the lack of enrichment for UCEs in loop anchors unsurprising. Indeed, unconstrained enhancers may more easily accommodate tissue-specific (108) or even species-specific regulatory programs (48).

To conclude, our data describe the pattern of relationships between ultraconservation of DNA sequence and three types of chromosome organization, with domains enriched in UCEs, boundaries being depleted, and loops being neither enriched nor depleted. More generally, they illustrate how different structural arenas of genome organization display distinct degrees of flexibility or stability over evolutionary timescales, as measured by ultraconservation.

## AVAILABILITY

Custom scripts associated with this study are available at http://github.com/rmccole/UCEsgenomeorganization.

## SUPPLEMENTARY DATA

Supplementary Data are available online. Supplementary Figures S1 to S5 are available as a single PDF file. Supplementary Tables S1 to S7 are available as separate excel files.

## ACKNOWLEDGMENTS

We thank Brian J. Beliveau, Chamith Y. Fonseka, Roxana Tarnita, Kaia Mattioli, Tommy Tullius, and all members of the Wu laboratory for valuable and insightful discussions.

## FUNDING

This work was supported by a William Randolph Hearst Foundation (to R.B.M.); an European Molecular Biology Organization (EMBO) Long-Term Fellowship [ALTF 186-2014 to J.E.]; National Institutes of Health (NIH) [DP1GM106412, R01HD091797 to C.-t.W.]; and Harvard Medical School (to C.-t.W.).

## Conflict of interest statement.

None declared.

